## Supplementary Info for "Local and substrate-specific S-palmitoylation determines subcellular localization of Gαo"

### Supplementary Methods

#### Screening of eukaryotic Gα subunits

To identify Gα subunit paralogs in eukaryotic genomes, the full-length amino acid sequence of Gαo was used for search queries in the NCBI protein Blast browser (blast.ncbi.nlm.nih.gov/Blast.cgi). Specifically, species with sequenced genomes were selected from Ensembl databases (ensembl.org) for vertebrates, plants, fungi, metazoa and protist. For the remaining phylogenetic clades, Gα paralogs were found as above and using the NCBI Taxonomy browser (ncbi.nlm.nih.gov/Taxonomy/Browser/wwwtax.cgi) as guide for clade classification. Sequences were manually curated and when annotated as partial were not considered further for this analysis. A total of 262 species were found to express Gα paralogs among the clades Amoebozoa, Discoba, Haptista, Metamonada, Opisthokonta, Sar, and Viridiplantae (Supplementary Data 1). Logo representation of the unique sequences selected for the different Gαo-$Nt^7$ categories (Supplementary Data 2) was done using the WebLogo application[1] (weblogo.berkeley.edu) and further edited with Inkscape v1.0.2 (inkscape.org).

### Supplementary Notes

#### The eukaryotic Gα-$Nt^7$

To determine how much diversity is present in $Nt^7$ among eukaryotic Gα subunits, we searched for Gα sequences containing an N-terminal heptapeptide with the signature $M^1$GXXX[ACGST]$X^7$, with at least one Cys at positions 3-5 or 7. Of note, we did not find any Gα subunit orthologue in the clades Ancyromonadida, Apusozoa, Breviatea, Cryptophyceae, CRuMs, Glaucocystophyceae, Malawimonadidae, and Rhodophyta (red algae). We found, however, Gα subunits in abundance in Amoebozoa, Discoba, Haptista, Metamonada, Opisthokonta, Sar, and Viridiplantae. Overall, we identified more than 800 sequences containing the desired $Nt^7$ signature (Supplementary Data 1), and uncovered some interesting features. For instance, no Gα subunits was found with a Cys occupying position 7. On the other hand, we observed Cys throughout positions 3-5, with Cys3 as the most common case with just over half of the sequences. Cys4 follows as the second most frequent with one third of all cases, whereas only ~6% of Gα subunits contain a Cys5. The vast majority of Gα subunits containing a Cys5 are from plants (~65%) and Amoebozoa (~25%). Curiously, Cys5 is not present in Fungi and is almost completely absent in Metazoa, with the genus *Caenorhabditis* as the sole identified exception (e.g. *C. elegans*; Supplementary Data 1). Less than 10% of all sequences present two Cys residues occupying positions 3-5, and they almost exclusively belong to fungal Gα subunits (Supplementary Data 1). A double Cys at positions 3 and 4 is by far the most frequent combination among them. No Gα sequence was found with a Cys occupying all three positions.

We asked whether consensus sequences might emerge from the Gα subunits among each category. As available databases are biased towards certain clades, we first selected unique hexapeptides with the signature M[1]GXXX[ACGST][6]. When an identical combination within positions 3-5 was found, we selected the sequence with the most frequent residue at position 7 for this analysis. Thus, we identified 88 unique combinations for Cys3, 75 for Cys4, and 13 for Cys5 (Supplementary Data 2). As seen in the corresponding Logo representations, there is no obvious Gα-Nt[7] consensus for the Cys3 and Cys4 categories (Notes Fig. 1a,b), whereas the sequence MGSLCSR emerges for Cys5 (Notes Fig. 1c). Plotting the amino acids by their polarity groups showed a common characteristic among categories: positively and even more so negatively charged amino acids are strongly underrepresented at positions 3-5 compared to a random distribution, or even completely absent as in the Cys5 category (Notes Fig. 1a-d). One can speculate that charged amino acids might interfere with at least one of the lipid modifications of Gα-Nt[7]. The most recent recognition motif for NMTs – M[1]G[^DEFRWY]X[^DEKR][ACGST][KR][7] – denotes that positive and negative residues are not favored at certain positions[2]. Accordingly, N-myristoylation of Gαo-Nt[7]-GFP appeared strongly impaired in N2a cells when different combinations of Arg, Asp, Glu, or Lys were introduced at positions 4 and 5 (Notes Fig. 1e). A Gαo-Nt[7]-GFP mutant with a His occupying position 5 showed a near normal membrane association regardless of the presence of a positive or negative residue at position 4 (Notes Fig. 1f). The dissimilar localization of these Gαo-Nt[7] mutants was not due to major variations in their expression levels (Notes Fig. 1g,h). Thus, N-myristoylation of Gα-Nt[7] of the Cys3 category appears optimal when Arg, Asp, Glu, or Lys residues are excluded from position 5, in total agreement with the substrate recognition motif described by Castrec *et al*[2].

**Notes Figure 1. Eukaryotic Gα-Nt7 sequences**

**a**-**d**, Logo representations (left) and quantification of frequency of amino acid polarity groups (right) of the unique Gα-Nt[7] sequences found in eukaryotic genomes of the categories Cys3 (**a**), Cys4 (**b**) and Cys5 (**c**). Frequency of a random distribution of amino acids (**d**). Amino acid polarity groups are color-coded as indicated in (**d**). **e**,**f**, Effect of positive and negative residues in the membrane association of Gαo-Nt[7]-GFP mutants inferred from their localization in N2a cells. Positions 4 and 5 (underlined) in Gαo-Nt[7] were point mutated to introduce negative (Asp or Glu) or positive (Arg, Lys or His) residues as indicated in the figures. Scale bars, 10 µm. **g**,**h**, Western blot analysis of N2a cells transfected as in **e** and **f**. Antibody against GFP was used to detect the expression level of Nt[7]-GFP constructs and α-Tubulin (α-Tub) as loading control (**g**). Quantification revealed no significant variation among the constructs (**h**). Data shown as the mean ± s.e.m. from 3 independent experiments. *P* values were determined using one-sample *t*-test for **h**. ns: not significant. Source data are provided as a Source Data file.

**Supplementary Figure 1. Gαo N-terminal region**

**a**-**c**, Overlay of the N-terminal region of three solved Gαo structures (**a**). N-terminus (**b**) and α-helix end (**c**) are magnified. Ala7 and Ala31 are underlined in Gαo sequence and color-coded in the structures (PDB-IDs): red for 6g79[3], blue for 6k41[4], yellow for 6oik[5]. The 6oik structure contained four substitutions indicated below the sequence in (**a**). **d**-**f**, Overlay of seven solved Gαi1 structures (**d**). Magnification of N-terminus (**e**) and α-helix end (**f**). Ala7 and Ala31 underlined in the sequence and color-coded in the structures (PDB-IDs): green for 5kdo[6], cyan for 1gg2[7], pink for 6ddf[8], yellow for 6k42[4], orange for 6kpf[9], red for 6n4b[10], blue for 6osa[11]. **g**,**h**, Expression in N2a cells of Gαo-GFP, Gαo-$Nt^{7}$-GFP or Gαo-$Nt^{31}$-GFP. Anti-GFP antibody was used to detect Gαo constructs and α-Tubulin (α-Tub) served as loading control (**g**). Quantification of Gαo constructs expression (**h**). Data as mean ± s.e.m.; 5 independent experiments. **i**, Localization of Gαo-$Nt^{8\text{-}31}$-GFP in N2a cells. **j**, Immunoprecipitation (IP) of Gαo-GFP, Gαo-$Nt^{7}$-GFP, Gαo-$Nt^{31}$-GFP or GFP as control. Only Gαo-GFP efficiently co-precipitated mRFP-Gβ1γ3. Antibodies against GFP and mRFP were used for detection of Gαo constructs and Gβ1γ3, respectively. **k**,**l**, Expressing of Gαo-$Nt^{7}$-GFP wild-type and mutants, and the consensus MGSLCSR-$Nt^{7}$-GFP (**k**). Underlined letters indicate residues mutated in Gαo-$Nt^{7}$. Detection and Quantification as in (**g**,**l**). Data as the mean ± s.e.m.; 3 independent experiments. **m**, [$^{3}$H]palmitate radiolabeling of HeLa cells expressing Gαo-$Nt^{7}$-GFP wild-type and mutants. Immunoprecipitated GFP-fusions from control (-) and hydroxylamine (+) treated samples were analyzed by autoradiography ([$^{3}$H]palmitate) and anti-GFP antibody. A long exposure is shown for visualization of weakly labelled constructs. **n**, Gαo-GFP localization in N2a cells treated with 2-bromopalmitate (2-BrPal). **o**-**q**, [$^{3}$H]palmitate radiolabeling of Gαo-GFP, MGNC-Gαo-GFP, and Gαo-C3N-GFP (**o**). GFP-fusions from control, 2-BrPal and Palmostatin B (PalmB) treated cells were analyzed as in (**m**). [$^{3}$H]palmitate incorporation to constructs normalized to corresponding control (**p**) or Gαo (**q**). Data as mean ± s.e.m.; 3 independent experiments. **i**,**n**, Scale bars, 10 µm. *P* values were determined using one-sample *t*-test for **h**,**l**,**p**,**q** and two-sided unpaired *t*-test for **q**. ns: not significant. Source data are provided as a Source Data file.

**Supplementary Figure 2. Cys position within Nt7 sequences**

**a**,**b**, Expression in N2a cells of Gαo-$Nt^{7}$-GFP wild-type and Ser6 mutants (**a**). Anti-GFP antibody was used to detect Gαo constructs and α-Tubulin (α-Tub) served as loading control. Quantification of $Nt^{7}$ constructs expression (**b**). Data as mean ± s.e.m.; 3 independent experiments. **c**, Localization of Ser6 mutants of Gαo-$Nt^{7}$-GFP. **d**,**e**, Co-expression of mRFP-

Lact-C2 with Gαo-Nt$^7$-GFP (top), MGNC-Nt$^7$-GFP (middle) or GalT-GFP (bottom) (**d**). DAPI stained nuclei in blue. Selected areas are magnified to the right. Underlined letters indicate residues substituted in Gαo-Nt$^7$. Pearson's co-localization analysis of GFP-fusions and mRFP-Lact-C2 (**e**). Box plots indicate median (middle line), 25th, 75th percentile (box), and lowest, highest value (whiskers); 3 independent experiments (Gαo-Nt$^7$, *n*=40; MGNC-Nt$^7$, *n*=39; GalT, *n*=42). **f**, Localization of MGNC-Nt$^7$-GFP upon 2-bromopalmitate (2-BrPal) incubation. **g**,**h**, Expression of GFP-fusions for full-length Gαo (Gαo-GFP), its MGNC mutant (MGNC-Gαo-GFP), or Gαo carrying the MGSLCSR sequence (MGSLCSR-Gαo-GFP). Immunodetection as in (**a**). Quantification of the expression of GFP-fusions (**h**). Data as mean ± s.e.m.; 3 independent experiments. **i**,**j**, MGNC-Gαo-GFP localization after 2-BrPal incubation (**i**) and MGSLCSR-Gαo-GFP under normal culture conditions (**j**). **k**, HeLa cells expressing Gαo-Nt$^7$-GFP (top), MGNC-Nt$^7$-GFP (middle), and MGSLCSR-Nt$^7$-GFP (bottom) were immunostained against GM130. Boxed areas are enlarged to the right. **l**-**o**, Live imaging of N2a cells co-expressing Gαo-Nt$^7$-GFP (**l**) or MGNC-Nt$^7$-GFP (**m**) and MannII-mRFP (bottom right insets). Representative images of the Nt$^7$-GFP constructs at the time of DMSO (Control) addition (0 min) and after 45 min. Quantification of PM content of Gαo-Nt$^7$ (**n**) and MGNC-Nt$^7$ (**o**) from Control and Palmostatin B (PalmB) treated cells in 5 min intervals and starting at t=0. Data represent mean ± s.d. (Gαo-Nt$^7$ control, *n*=20; Gαo-Nt$^7$ PalmB, *n*=20; MGNC-Nt$^7$ control, *n*=20; MGNC-Nt$^7$ PalmB, *n*=20). **p**,**q**, Localization of Gαo-GFP (**p**) and MGNC-Gαo-GFP (**q**) in N2a cells after 45 min of PalmB treatment. **c**,**d**,**f**,**i**-**m**,**p**,**q**, Scale bars, 10 µm. *P* values were determined using one-sample *t*-test for **b**,**h** and one-way ANOVA Tukey test for **e**. ns: not significant. Source data are provided as a Source Data file.

**Supplementary Figure 3. Subcellular localization of zDHHCs in N2a cells**

**a**, A protein-lipid overlay assay on membrane strips spotted with different lipids as indicated in the illustration (left). Membrane strips were incubated with extracts from N2a cells expressing the control FAPP1-PH-GFP, Gαo-Nt$^7$-GFP, MGNC-Nt$^7$-GFP or Gαo-Nt$^{31}$-GFP, and detection of the constructs was done using an Ab against GFP. Images of the membrane strips are shown side-by-side with the corresponding Ab signals. **b**, N2a cells expressing the HA-tagged zDHHC collection were immunostained against HA and GM130 (not shown) to determine the subcellular localization of zDHHCs. Scale bars, 10 µm.

**Supplementary Figure 4. Validation of the S-palmitoylation at the ONM – SwissKASH – assay**

**a**-**c**, The SwissKASH assay in HeLa cells applied to GFP-fusions of SNAP23 (GFP-SNAP23; **a**), caveolin-1 (GFP-Cav1; **b**), and flotllin-2 (Flot2-GFP; **c**). GFP-SNAP23 was efficiently recruited to the ONM by the KASH-constructs of zDHHC13 (mRFP-zDHHC13-KASH) and zDHHC17 (mRFP-zDHHC17-KASH), but not by the control mRFP-KASH (**a**). Similarly, GFP-Cav1 targeted the ONM by the co-expression of mRFP-zDHHC7-KASH (**b**), and Flot2-GFP by mRFP-zDHHC5-KASH (**c**), but not by the control mRFP-KASH (**b**,**c**). **d**,**e**, The SwissKASH assay for Gαo-Nt$^7$-GFP (**d**) and MGNC-Nt$^7$-GFP (**e**) using the mRFP- and KASH-fusions of the PM-associated zDHHC2, 5, 8, 14, and 18. Underlined letters indicate residues substituted in Gαo-Nt$^7$. **a**-**e**, DAPI staining of nuclei in blue, boxed areas are enlarged to the right. Scale bars, 10 µm.

**Supplementary Figure 5. Golgi-localized zDHHCs**

**a**,**b**, The SwissKASH assay in HeLa cells applied to Gαo-Nt$^7$-GFP (**a**) and MGNC-Nt$^7$-GFP (**b**) using the mRFP- and KASH-fusions of the Golgi-associated zDHHC9, 12, 13, 15, 16, 17, 21, 23 and 25. Underlined letters indicate residues substituted in Gαo-Nt$^7$. A blue DAPI staining marks nuclei. Selected areas are zoomed-in to the right. Scale bars, 10 µm.

**Supplementary Figure 6. Substrate specificity of zDHHC11**

**a**-**c**, The SwissKASH assay in HeLa cells was applied to MGSSCSR-Nt$^7$-GFP (**a**) and MGLLCSR-Nt$^7$-GFP (**b**) using the control mRFP-KASH or mRFP-zDHHC11-KASH constructs. Marked areas are magnified to the right. Quantification of mean fluorescence intensity ratio of GFP-constructs at the ONM versus cytosol (**c**). Box plots indicate median (middle line), 25th, 75th percentile (box), and lowest, highest value (whiskers) from 3 independent experiments (MGSSCSR-Nt$^7$, $n$=49; MGLLCSR-Nt$^7$, $n$=49). **d**-**g**, Immunoprecipitation (IP) of SUN2-GFP, Gαo-Nt$^7$-GFP and MGNC-Nt$^7$-GFP (**d**-**f**) or MGSSCSR-Nt$^7$-GFP and MGLLCSR-Nt$^7$-GFP (**g**) under the co-expression of mRFP-zDHHC3-KASH (**d**), mRFP-zDHHC7-KASH (**e**) or mRFP-zDHHC11-KASH (**f**,**g**). Underlined letters indicate residues substituted in Gαo-Nt$^7$. Note that only SUN2-GFP co-precipitated all mRFP-zDHHC-KASH constructs. Antibodies against GFP and mRFP were used for detection. **a**,**b**, Scale bars, 10 µm. $P$ values were determined using two-sided unpaired $t$-test for **c**. Source data are provided as a Source Data file.

**Supplementary Figure 7. Accessory proteins in the SwissKASH assay**

**a**-**d**, The SwissKASH assay for Gαo-Nt7-GFP (**a**,**c**) and MGNC-Nt7-GFP (**b**,**d**) using mRFP-zDHHC5-KASH (**a**,**b**) or mRFP-zDHHC9-KASH (**c**,**d**), both in conjunction with GCP16/Golga7-Flag. Cells were immunostained against Flag-tag and nuclei were stained in blue with DAPI. Marked regions are magnified at the bottom panels. Underlined letters indicate residues substituted in Gαo-Nt$^7$. **e**-**g**, The SwissKASH assay applied for

GCP16/Golga7-Flag using the control mRFP-KASH (**e**), mRFP-zDHHC5-KASH (**f**) or its DHHS-inactive mutant (mRFP-zDHHS5-KASH; **g**). Note that the strong ONM accumulation of GCP16/Golga7-Flag induced by the KASH-fusion of zDHHC5 is not seen by its DHHS mutant. Cells were immunostained against Flag-tag and DAPI stained nuclei in blue. Selected areas are enlarged to the right. **h**, Immunoprecipitation (IP) of SUN2-GFP, Gαo-Nt$^{7}$-GFP and MGNC-Nt$^{7}$-GFP by the co-expression of mRFP-zDHHC5-KASH and Golga7b-Flag. Note that only SUN2-GFP co-precipitated mRFP-zDHHC5-KASH. Golga7b was weakly detected in the input and not detected at all in the IP (no shown). Antibodies (Abs) against GFP, mRFP, and Flag-tag were used for detection. **i**, The SwissKASH assay in HeLa cells was applied to Gαo-Nt$^{31}$-GFP using mRFP-zDHHC5-KASH together with Golga7b-Flag. Ab against Flag-tag was used for immunostaining and DAPI for staining nuclei in blue. A marked area is magnified at the bottom. **a**-**g**,**i**, Scale bars, 10 µm. Source data are provided as a Source Data file.

**Supplementary Figure 8. Localization and expression of Nt$^{7}$ constructs upon co-expression of zDHHCs**

**a**,**b**, Representative images of N2a cells expressing the Golgi marker MannII-BFP together with Gαo-Nt$^{7}$-GFP (**a**) or MGNC-Nt$^{7}$-GFP (**b**). Selected areas are zoomed-in to the right. Underlined letters indicate residues substituted in Gαo-Nt$^{7}$. **c**-**f**, Western blot (WB) analysis of N2a cells. Cells were co-transfected with Gαo-Nt$^{7}$-GFP together with empty pcDNA3.1(+) plasmid (Control), HA-tagged zDHHC11 (HA-zDHHC11) or its DHHS-inactive mutant (HA-zDHHS11) (**c**). MGNC-Nt$^{7}$-GFP was co-transfected with control plasmid, HA-zDHHC3 or HA-zDHHS3 (**d**), as well as with HA-zDHHC7 or HA-zDHHS7 (**e**). Antibody (Ab) against GFP was used to detect the expression level of Nt$^{7}$ constructs, anti-HA detected HA-zDHHCs and α-Tubulin (α-Tub) served as loading control (**c**-**e**). Arrowheads marked as Nt$^{7}$ point to anti-GFP residual signals. Quantification showed no significant (ns) difference between the Nt$^{7}$-GFP constructs (**f**). Data shown as the mean ± s.e.m. from 3 independent experiments. **g**-**i**, Confocal images of N2a cells expressing Gαo-Nt$^{7}$-GFP (**g**,**h**) or MGNC-Nt$^{7}$-GFP (**i**) together with HA-zDHHC3 (**g**), HA-zDHHC7 (**h**), or HA-zDHHC11 with MannII-BFP (**i**). Cells were immunostained against the HA-tag. DAPI staining was used to visualize nuclei (**g**,**h**). **j**,**k**, Co-expression of Gαo-Nt$^{7}$-GFP (**j**) or MGNC-Nt$^{7}$-GFP (**k**) with HA-zDHHC5 and Golga7b-Flag in N2a cells. Immunostainings against the HA- and Flag-tags were used for detection and DAPI staining to mark nuclei in blue. **a**,**b**,**g**-**k**, Scale bars, 10 µm. *P* values were determined using one-sample *t*-test for **f**. ns: not significant. Source data are provided as a Source Data file.

**Supplementary Figure 9. Full-length Gαo and the ER-localized zDHHCs**

**a**-**c**, Confocal images of HeLa cells co-expressing HA-zDHHC3 (**k**), HA-zDHHC7 (**l**), or HA-DHHC11 (**m**) with the Golgi marker MannII-GFP. Cells were immunostained against the HA-tag, and DAPI stained in blue the nuclei. **d**, The SwissKASH assay for the MGNC mutant of full-length Gαo (MGNC-Gαo-GFP) using the mRFP-KASH control, zDHHC3, 7 and 11. Note that MGNC-Gαo-GFP efficiently targeted the ONM by co-expression of zDHHC11, but not of zDHHC3 and 7. Underlined letters indicate residues substituted in Gαo. DAPI staining of nuclei in blue. Selected regions are zoomed-in to the right. **e**, Representative images of HeLa cells co-expressing the GFP-fusion of the full-length Gαo (Gαo-GFP) together with the HA-tagged zDHHCs that localize at the ER (HA-zDHHC1, 4, 6, 19, 20 and 24). Cells were immunostained against the HA-tag and nuclei were stained in blue with DAPI. White arrowheads point to zDHHC1 and 20 at filopodia, indicating their localization at the PM. Note that co-expression of the ER-localized zDHHCs seemed not to affect the overall localization of Gαo-GFP. **a**-**e**, Scale bars, 10 µm.

**Supplementary Figure 10. The SwissKASH assay carrying zDHHC1 and 20**

**a**-**d**, The SwissKASH assay was applied to Gαo-$Nt^7$-GFP (**a**,**c**) and MGNC-$Nt^7$-GFP (**b**,**d**) using mRFP-zDHHC1-KASH (**a**,**b**) or mRFP-zDHHC20-KASH (**c**,**d**). Underlined letters indicate residues substituted in Gαo-$Nt^7$. A blue DAPI staining marks nuclei. Marked regions are magnified to the right. **e**, Localization of full-length Gαo-GFP and mRFP-Gβ1γ3 in HeLa cells co-expressing the SwissKASH control construct BFP-KASH (not shown). **f**, The SwissKASH assay applied to Gαo-GFP and the $His_6$-tagged RGS19 (His-RGS19) using BFP-zDHHC11-KASH (not shown). Cells were immunostained against the $His_6$-tag. A selected area is enlarged at the bottom. **g**,**h**, Quantification of the apparent PM and Golgi localization of Gαo-$Nt^7$-GFP and Gαo-$Nt^{31}$-GFP from images recorded using the same confocal setting. A scatter plot (**g**) shows that to a similar Golgi content of both constructs, Gαo-$Nt^{31}$ showed higher PM values (arbitrary (arb.) units). Dashed lines represent linear regressions with their equations alongside. A bar graph (**h**) of the pooled data illustrates the variation between the relative Golgi and PM content of both constructs. Box plots indicate median (middle line), 25th, 75th percentile (box), and lowest, highest value (whiskers) from 2 independent experiments (Gαo-$Nt^7$, $n$=56; Gαo-$Nt^{31}$, $n$=57). **a**-**f**, Scale bars, 10 µm. $P$ values were determined using two-sided unpaired $t$-test for **h**. Source data are provided as a Source Data file.

**Supplementary Data 1.**

Eukaryotic species and their Gα subunit sequences (N-terminal 10 amino acids) with the corresponding Entrez IDs. Phylogenetic clade classification was according to the NCBI Taxonomy browser.

**Supplementary Data 2.**

The unique Gα-$Nt^7$ sequences (in fasta format) found in eukaryotic genomes of the categories Cys3, Cys4 and Cys5.

**Supplementary Data 3.**

Sequences of oligonucleotides used in this study.

**Supplementary Movie 1.**

N2a cells expressing Gαo-$Nt^7$-GFP and the Golgi marker MannII-mRFP were treated with 50 µM Palmostatin B, and immediately recorded at one image per 30 seconds for 45 min as described in Methods. The Movie at 10 frames per second shows several cells displaying the changes in localization of Gαo-$Nt^7$ upon time.

**Supplementary Movie 2.**

N2a cells expressing MGNC-$Nt^7$-GFP and the Golgi marker MannII-mRFP were treated and recorded as in Supplementary Movie 1. The Movie shows a group of cells displaying the changes in MGNC-$Nt^7$ localization upon time.

**Supplementary Movie 3.**

HeLa cells expressing Gαo-$Nt^7$-FM4-GFP and the Golgi marker MannII-mRFP were treated with D/D solubilizer, and immediately recorded at one image per 5 seconds for 10 min as described in Methods. The Movie at 12 frames per second shows two representative cells displaying the Golgi accumulation of Gαo-$Nt^7$-FM4-GFP upon time.

**Supplementary Movie 4.**

HeLa cells expressing MGNC-$Nt^7$-FM4-GFP and MannII-mRFP were prepared and recorded as in Supplementary Movie 3. The Movie shows two representative cells showing a mainly PM targeting of MGNC-$Nt^7$-FM4-GFP.
