## Supplementary figures and images for "Local and substrate-specific S-palmitoylation determines subcellular localization of Gαo"

### Notes Figure 1

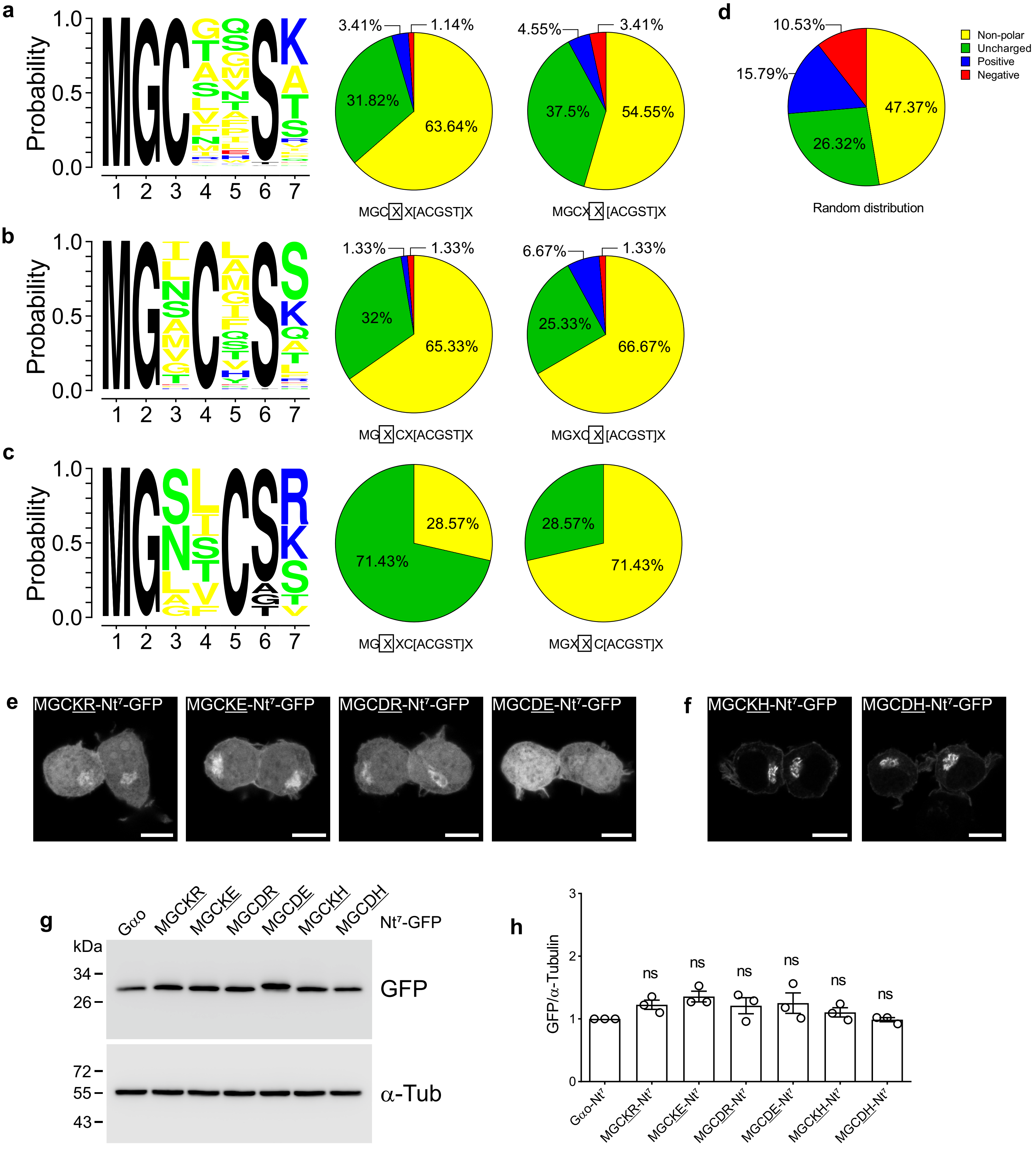

### Supplementary Figure 1

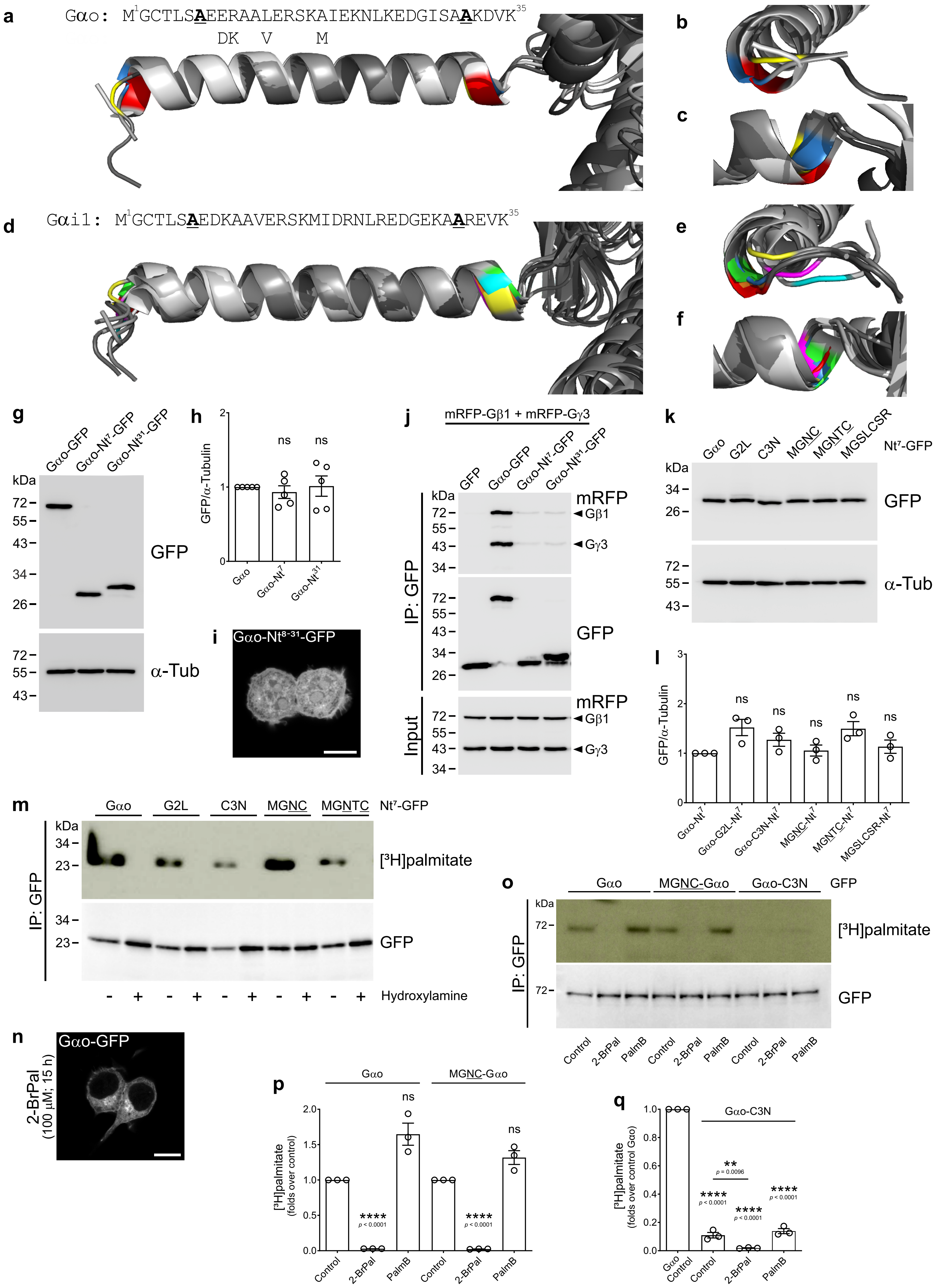

### Supplementary Figure 2

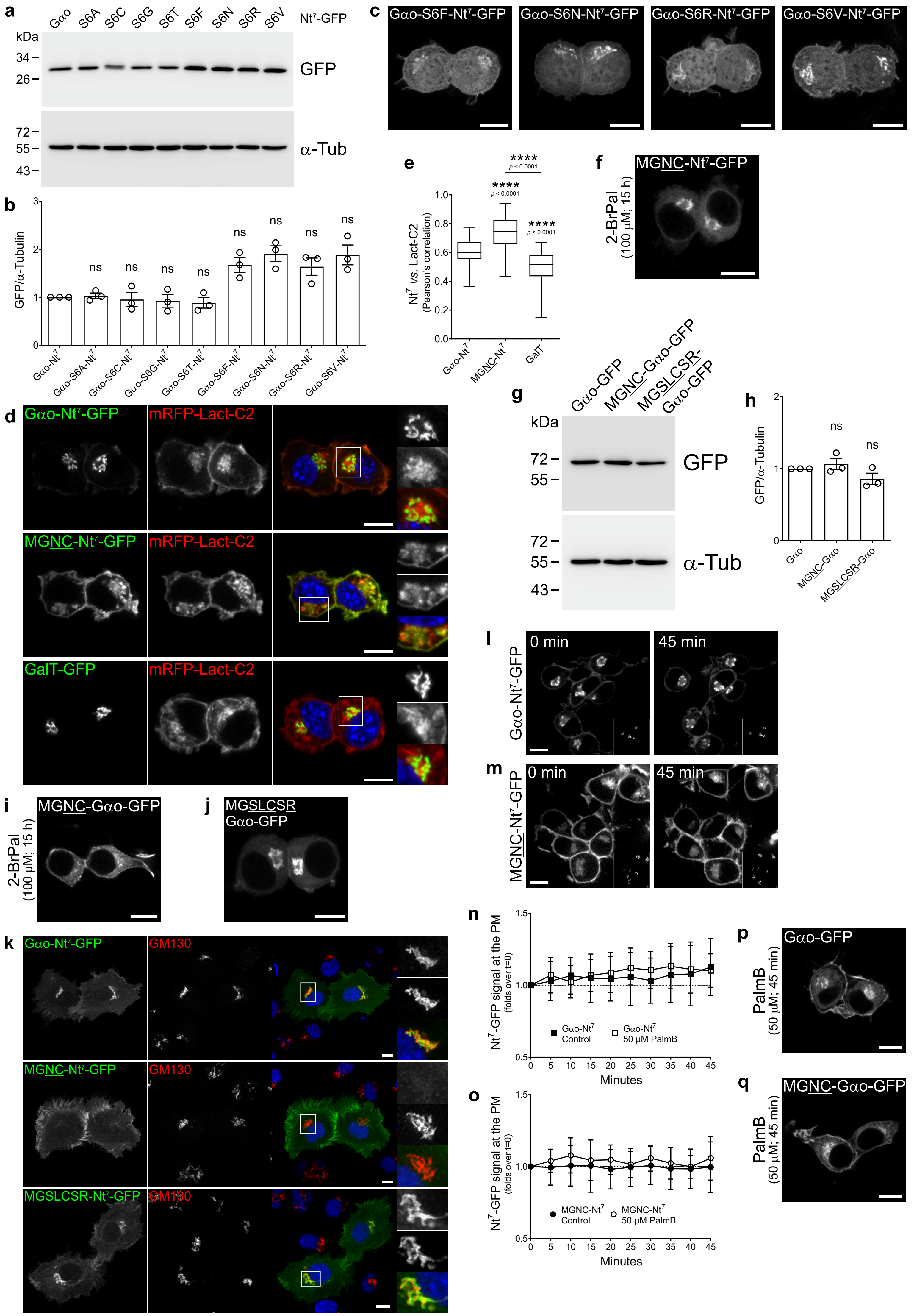

### Supplementary Figure 3

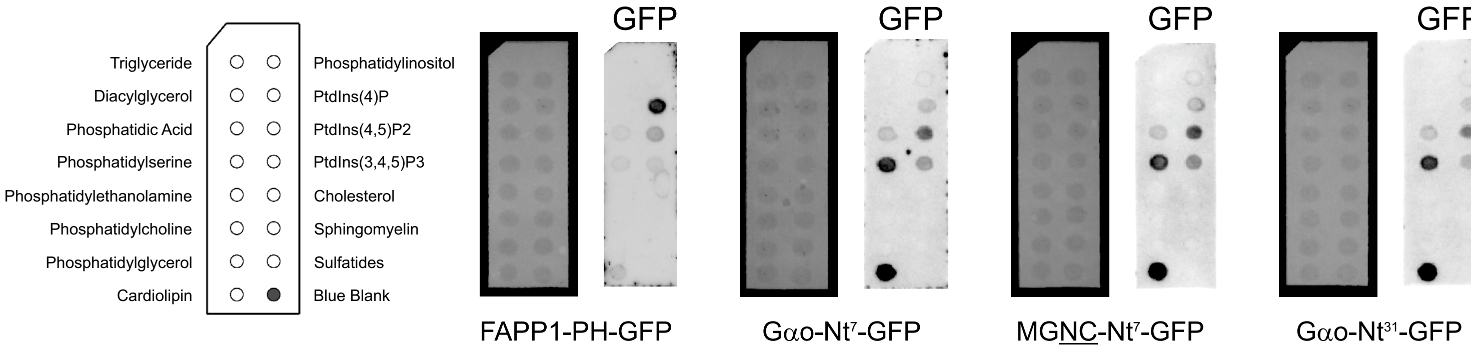

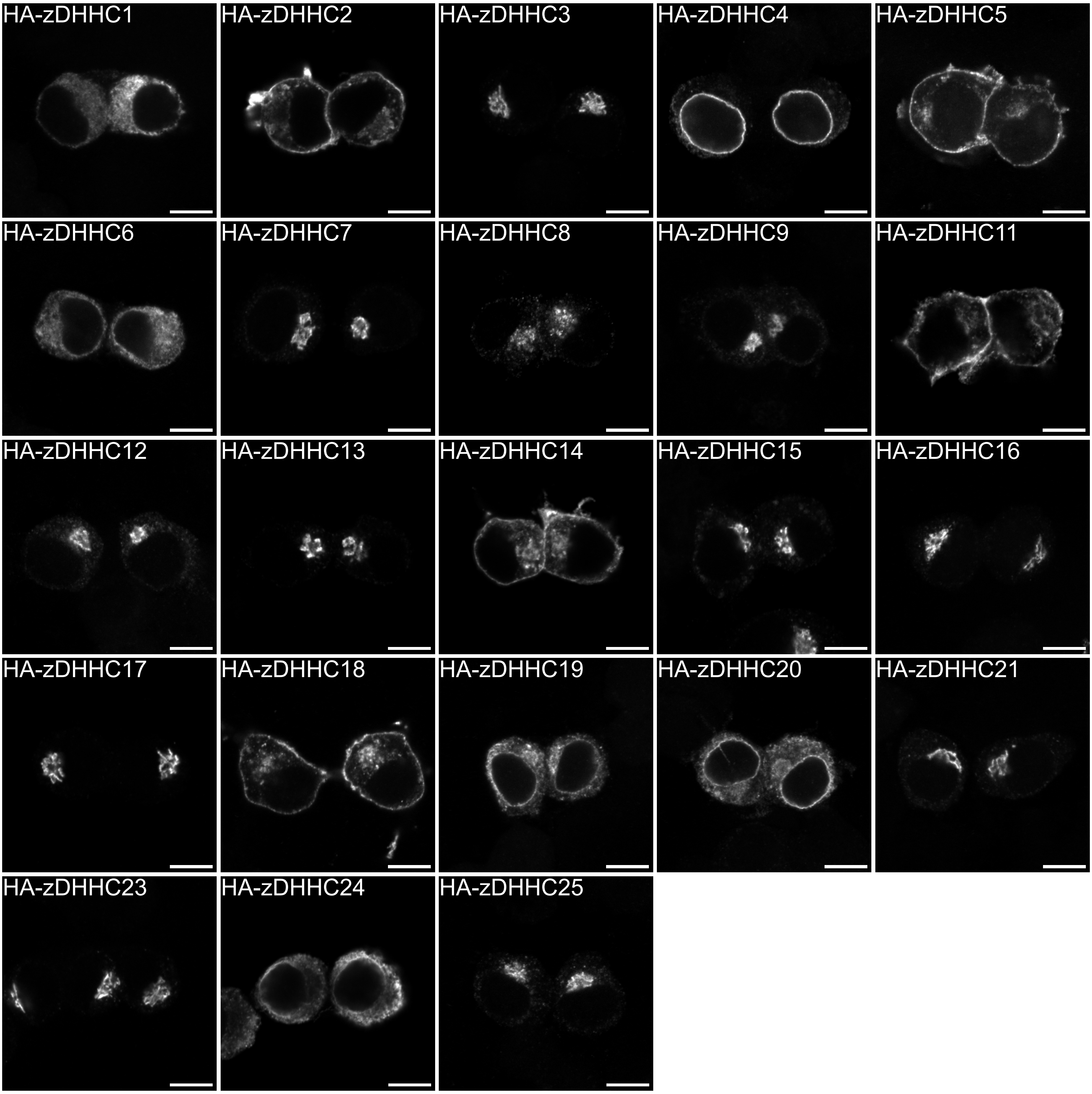

### Supplementary Figure 4

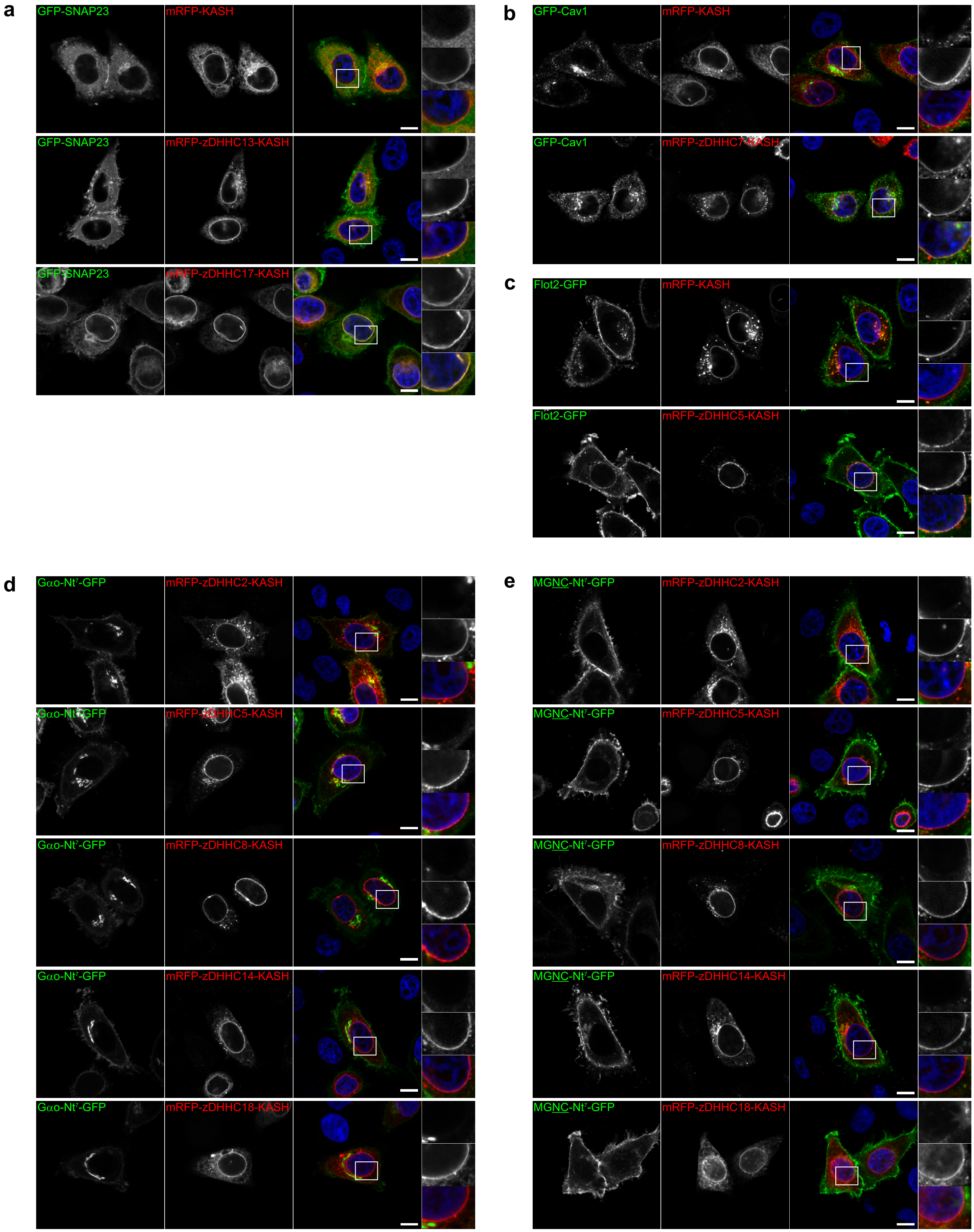

### Supplementary Figure 5

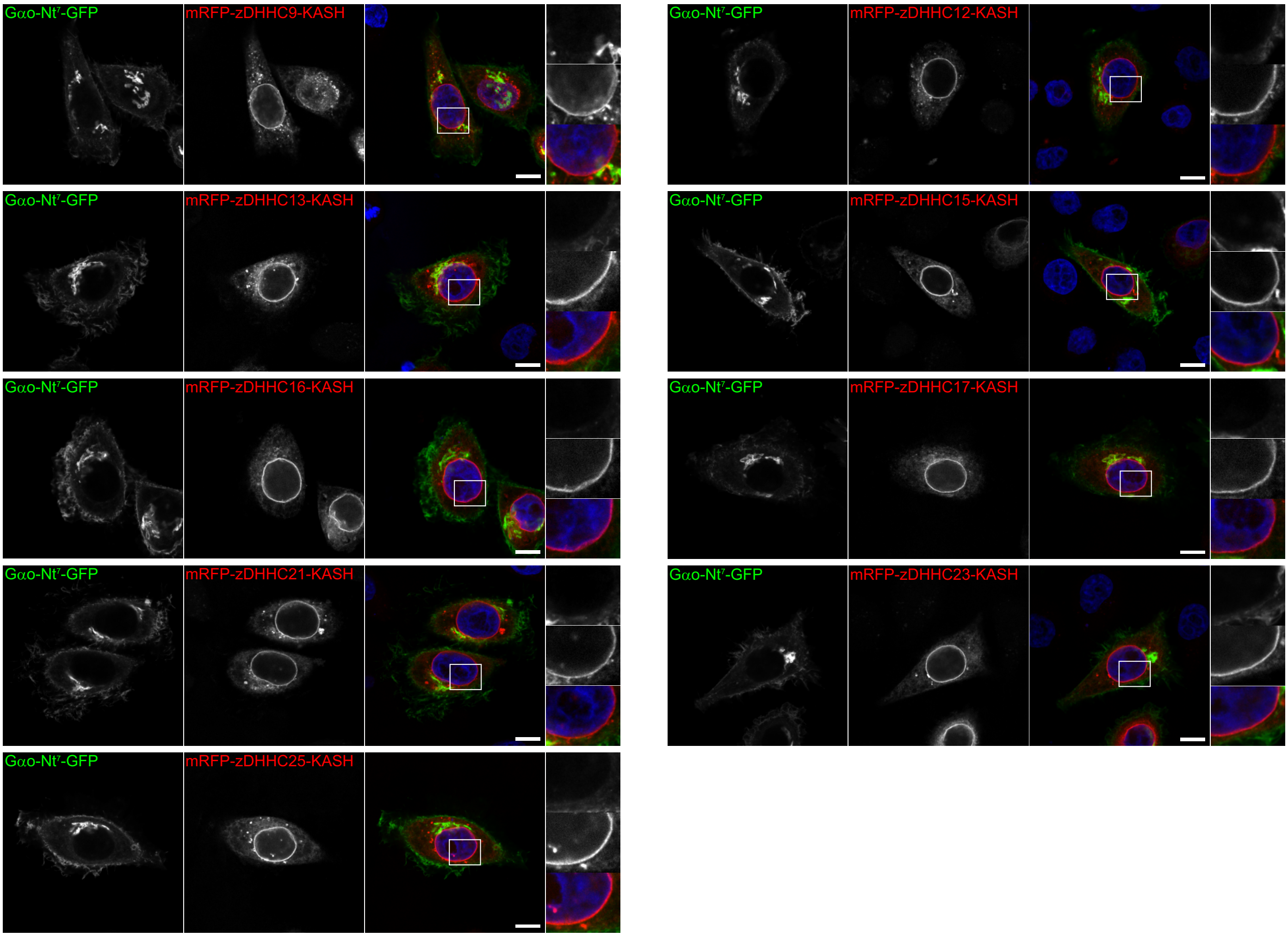

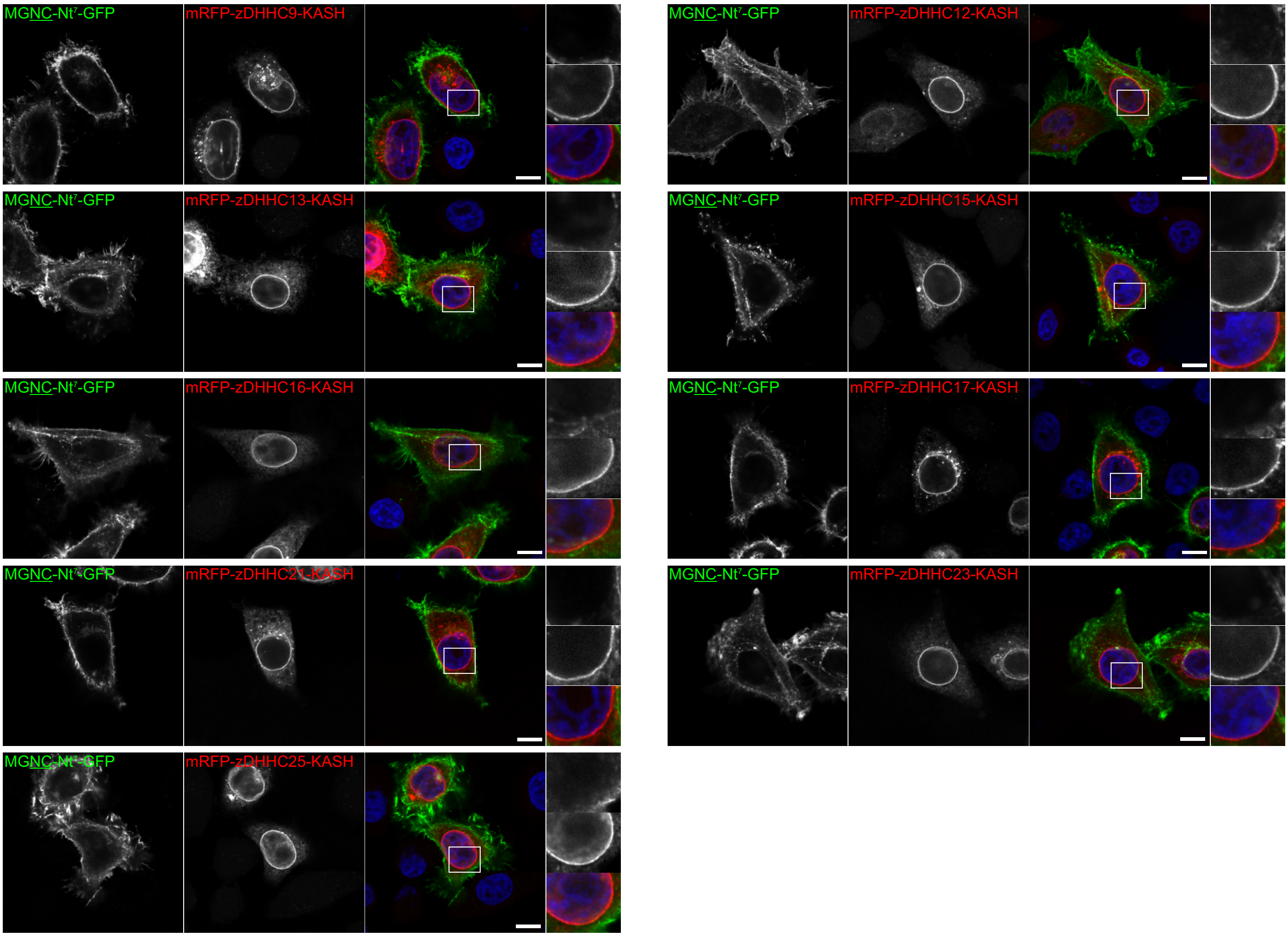

### Supplementary Figure 6

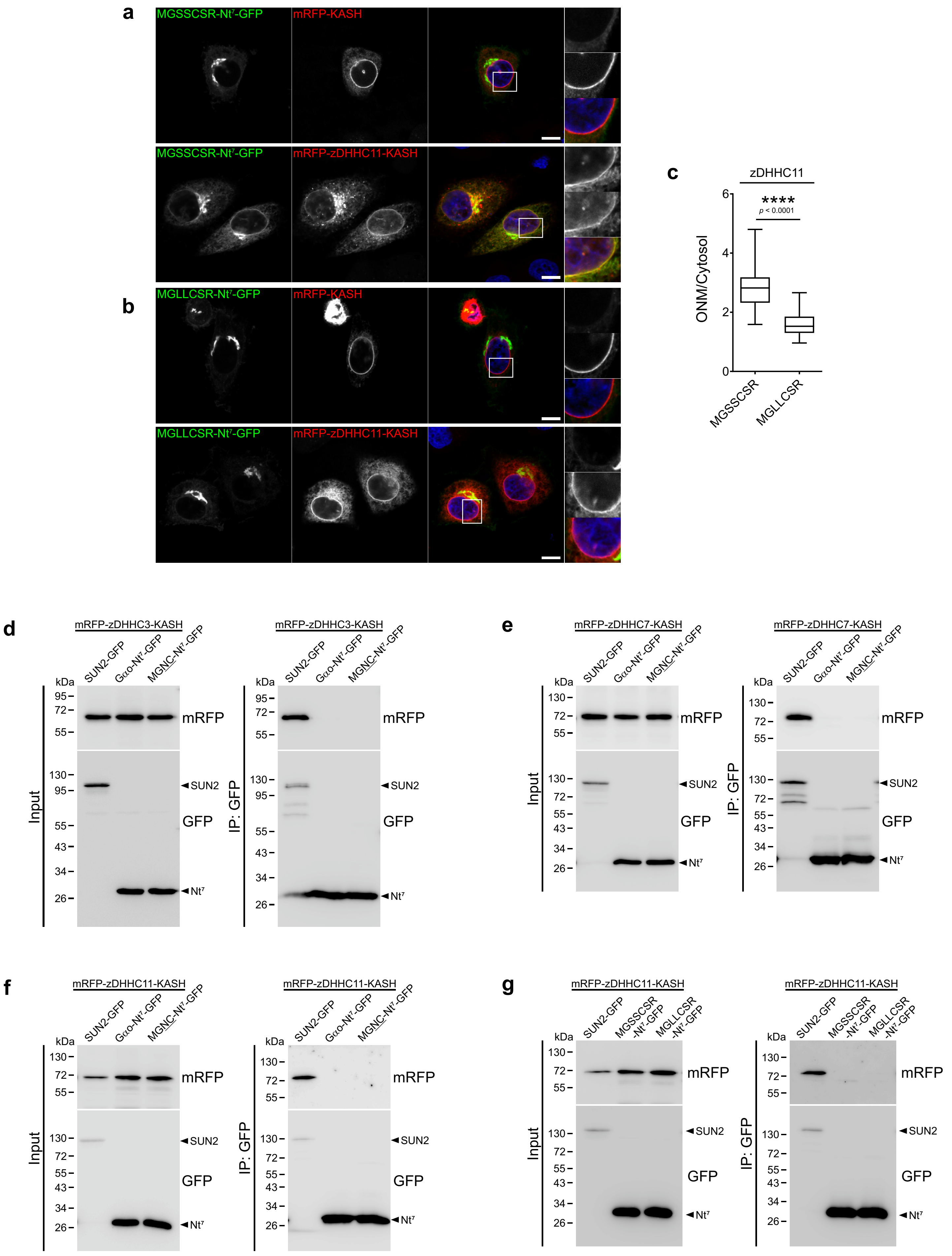

### Supplementary Figure 7

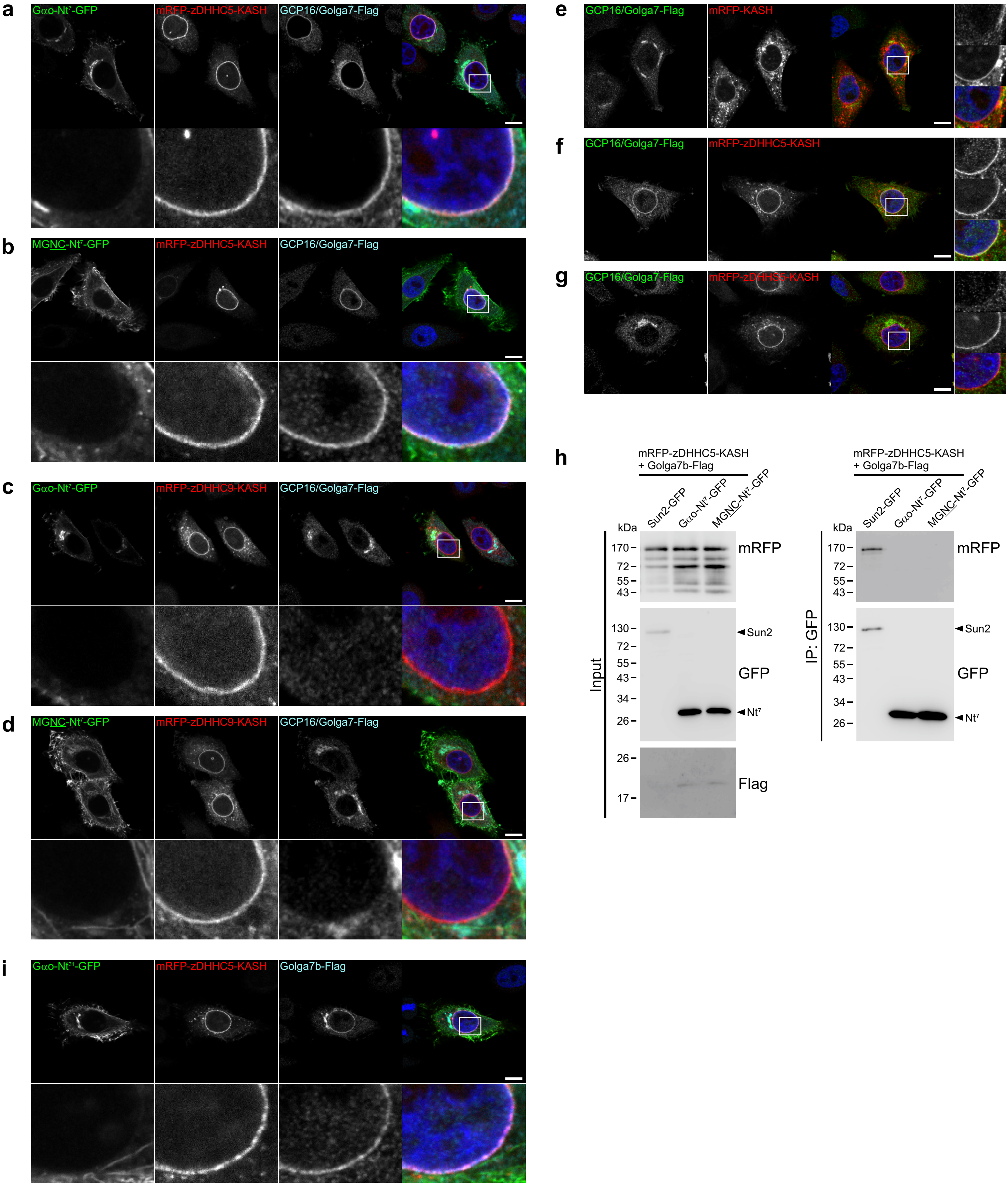

### Supplementary Figure 8

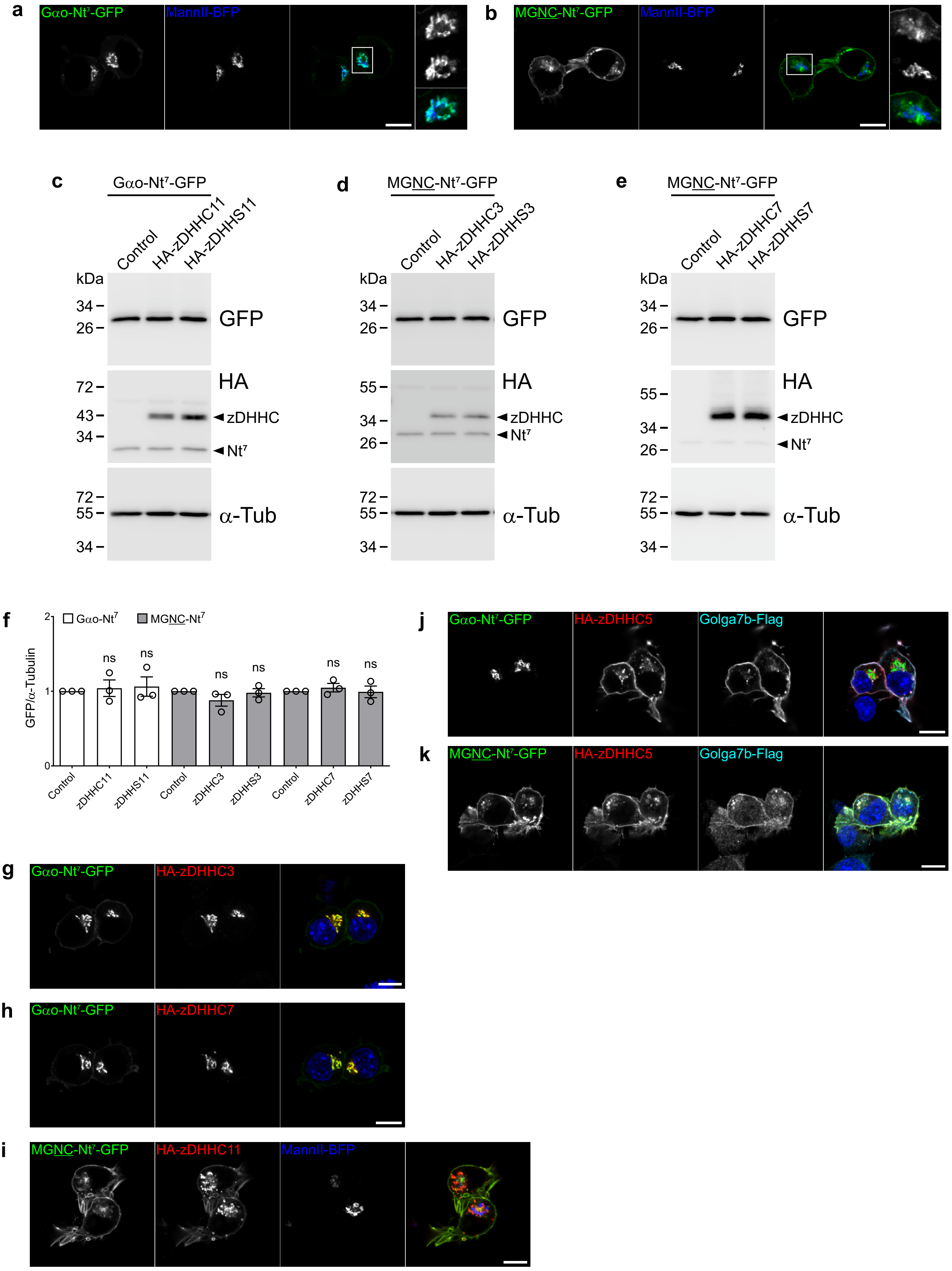

### Supplementary Figure 9

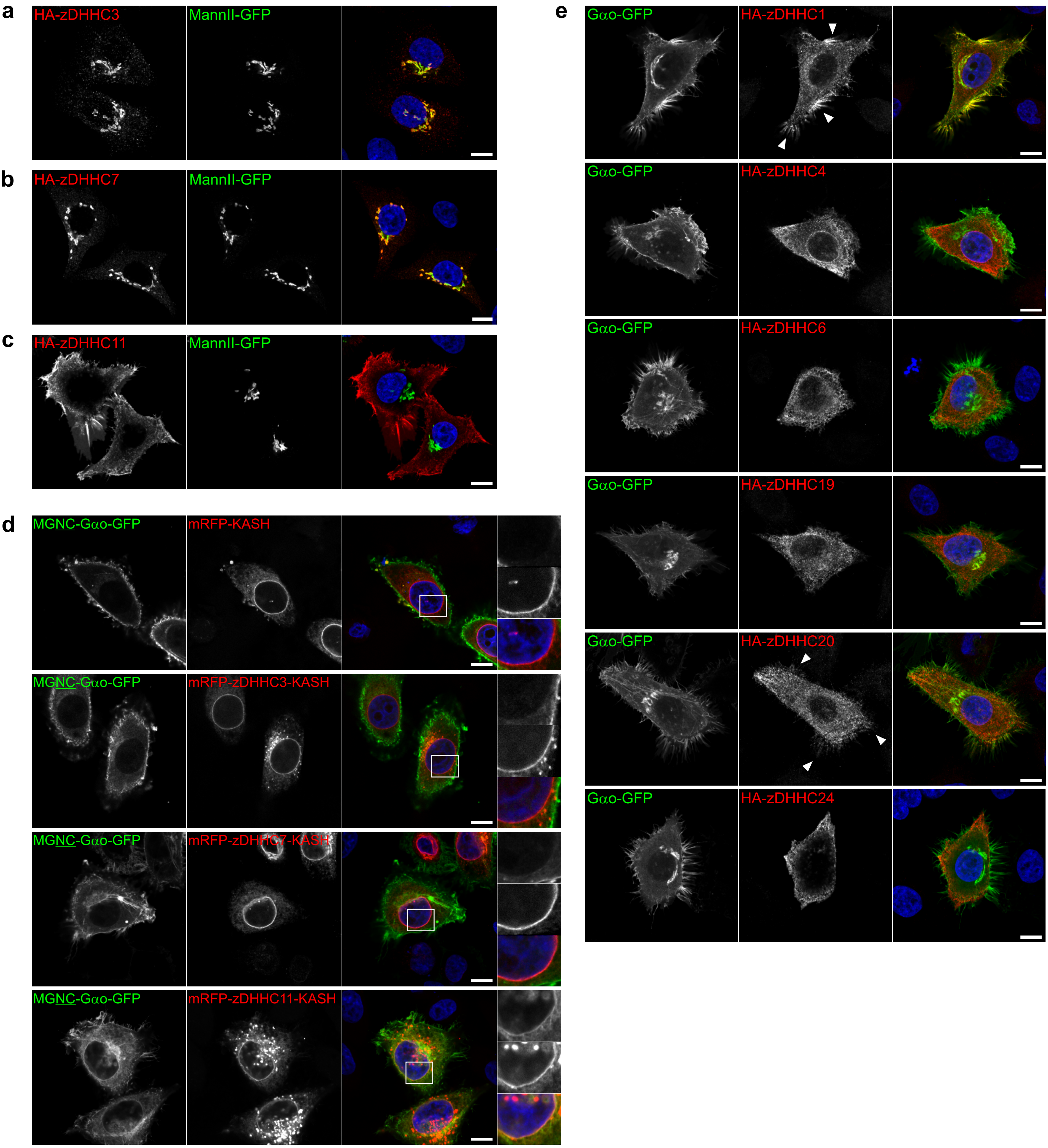

### Supplementary Figure 10

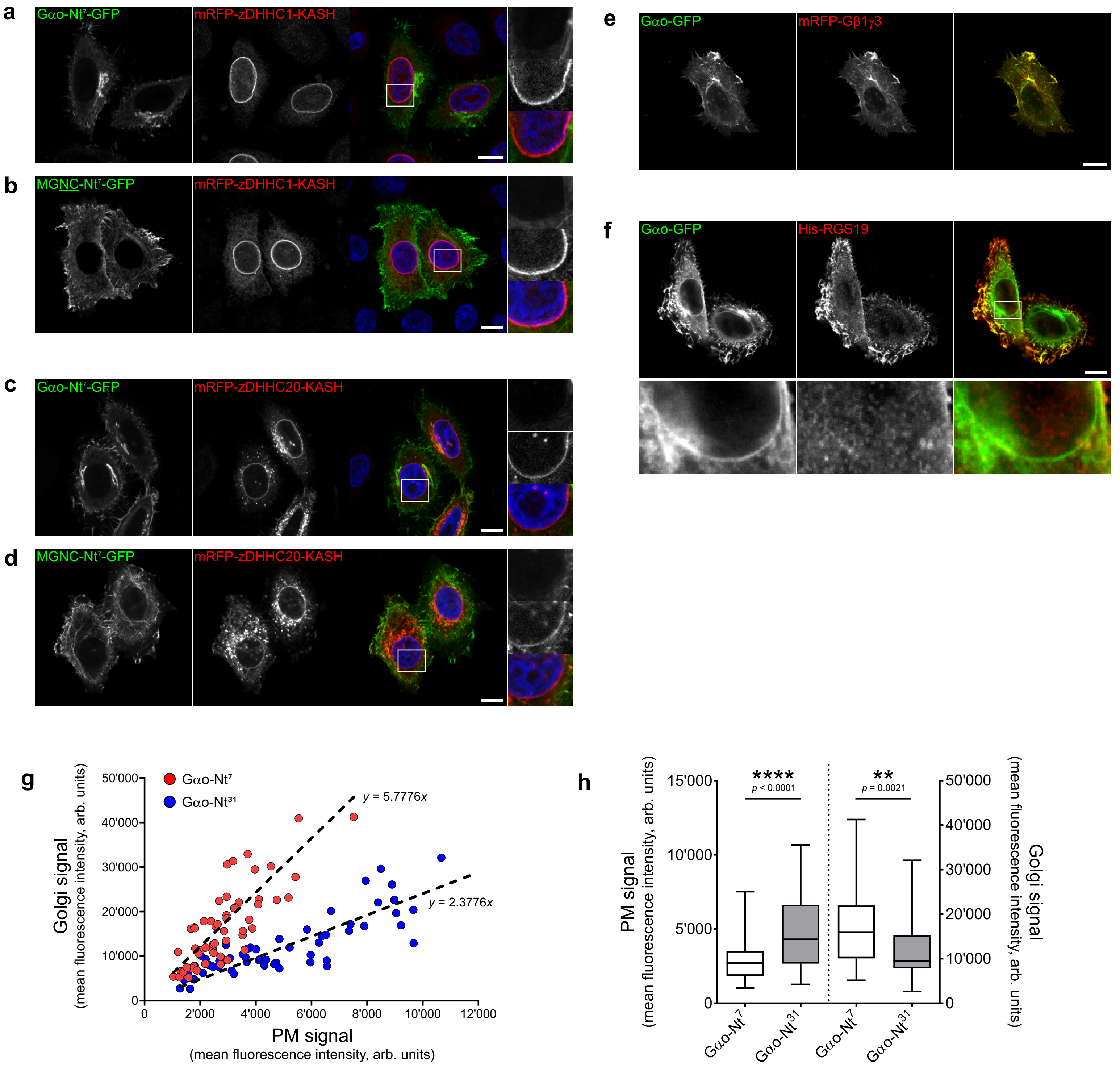
